# A musculoskeletal finite element model of rat knee joint for evaluating cartilage biomechanics during gait

**DOI:** 10.1101/2021.09.01.458496

**Authors:** Gustavo A. Orozco, Kalle Karjalainen, Eng Kuan Moo, Lauri Stenroth, Petri Tanska, Jaqueline Lourdes Rios, Teemu V. Tuomainen, Mikko J. Nissi, Hanna Isaksson, Walter Herzog, Rami K. Korhonen

## Abstract

Abnormal loading of the knee due to injuries or obesity is thought to contribute to the development of osteoarthritis (OA). Small animal models have been used for studying OA progression mechanisms. However, numerical models to study cartilage responses under dynamic loading in preclinical animal models have not been developed. Here we present a musculoskeletal finite element (FE) model of a rat knee joint to evaluate cartilage biomechanical responses during a gait cycle. The rat knee joint geometries were obtained from a 3-D MRI dataset and the boundary conditions regarding loading in the joint were extracted from a musculoskeletal model of the rat hindlimb. The fibril-reinforced poroelastic (FRPE) properties of the rat cartilage were derived from data of mechanical indentation tests. Our numerical results showed the relevance of simulating anatomical and locomotion characteristics in the rat knee joint for estimating tissue responses such as contact pressures, stresses, strains, and fluid pressures. We found that the contact pressure and maximum principal strain were virtually constant in the medial compartment whereas they showed the highest values at the beginning of the gait cycle in the lateral compartment. Furthermore, we found that the maximum principal stress increased during the stance phase of gait, with the greatest values at midstance. We anticipate that our approach serves as a first step towards investigating the effects of gait abnormalities on the adaptation and degeneration of rat knee joint tissues and could be used to evaluate biomechanically-driven mechanisms of the progression of OA as a consequence of joint injury or obesity.

**Author Summary:** Osteoarthritis is a disease of the musculoskeletal system which is characterized by the degradation of articular cartilage. Changes in the knee loading after injuries or obesity contribute to the development of cartilage degeneration. Since injured cartilage cannot be reversed back to intact conditions, small animal models have been widely used for investigating osteoarthritis progression mechanisms. Moreover, experimental studies have been complemented with numerical models to overcome inherent limitations such as cost, difficulties to obtain accurate measures and replicate degenerative situations in the knee joint. However, computational models to study articular cartilage responses under dynamic loading in small animal models have not been developed. Thus, here we present a musculoskeletal finite element model of a rat knee joint to evaluate cartilage biomechanical responses during gait. Our computational model considers both the anatomical and locomotion characteristics of the rat knee joint for estimating mechanical responses in the articular cartilage. We suggest that our approach can be used to investigate tissue adaptations based on the mechanobiological responses of the cartilage to prevent the progression of osteoarthritis.

## Introduction

Abnormal loading of the knee joint after overuse, severe injuries, or obesity are risk factors of cartilage degeneration, contributing to the development of osteoarthritis (OA) [1]. OA is the most common musculoskeletal disorder and among the most frequent causes of pain, physical disability, and economic loss worldwide [2]. Currently, there is no cure for OA, and patients with end-stage OA must undergo a total joint replacement to recover mobility and relieve the pain. Although it is understood that the mechanical environment plays a role in the onset and development of OA, the mechanisms leading to the progression of OA remain largely unknown, thereby preventing the development of effective measures to stop or slow down the degeneration of the joint [3,4].

In order to comprehend the degenerative mechanisms, preclinical animal models have been used in orthopaedic research for studying the initiation and progression of OA [5–7]. In preclinical research, small animal models (e.g., rodents) are commonly used as they are cost-effective and take less time to respond to an intervention compared to large animal models [8]. Invasive and non-invasive models have been developed to study different OA phenotypes. For example, invasive models utilize surgical injuries (ACL transection, meniscectomy, and destabilization of medial meniscus (DMM)) or chemical interventions to induce cartilage degradation (intra-articular injections of proinflammatory cytokines) [9,10]. On the other hand, noninvasive models include load-induced impact injury, cyclic joint loading, or spontaneous/genetic OA development [11–13].

Experimental studies have been complemented with numerical models to overcome inherent limitations such as cost, challenges to obtain accurate measures experimentally *in vivo*, and replicate degenerative scenarios in the knee joint. Finite element (FE) models have been used to investigate knee joint function during locomotion and joint loading alterations, as well as the associated adaptation and degeneration in the joint tissues [14,15]. For instance, subject-specific FE models of the knee joint have been developed to study the biomechanical responses of articular cartilage and meniscus after ACL rupture and reconstruction [16,17]. These computational models include realistic knee tissue geometries acquired from magnetic resonance imaging (MRI) data, complex material models to account for tissue anisotropy, and dynamic loading from a patient’s gait or other relevant motion, to provide insights into the role of biomechanics in the development of OA. Since physiological changes in articular cartilage occur faster in rodents, these realistic numerical models would be helpful to investigate the effect of treatments on cartilage tissue. Nevertheless, only a few simplified FE models for joints of rodents have been reported in the literature [18,19]. In previous studies, micro-computed tomography (μCT) imaging was used to obtain the geometry of the cartilages, bone, and meniscus that were subsequently implemented in FE models [20,21]. However, those studies assumed the cartilage thickness based on the proximal tibia and distal femur segmentations, simulated simplified loading conditions in the numerical model (e.g., only standing posture), and adopted cartilage tissue to be isotropic and linearly elastic, limiting the use of these models in preclinical rodent studies of OA.

In order to use FE modeling to understand mechanisms leading to OA in animal models, a methodology has to be developed first. In this study, we developed an FE model of a rat knee joint to estimate articular cartilage biomechanics during the stance phase of gait. The FE model included a fibril-reinforced poroelastic (FRPE) material model that accounts for material nonlinearities of meniscus and cartilages, as well as their nonfibrillar and fibrillar matrices. The FRPE properties of the rat cartilage were obtained by fitting the model to previous indentation experiments [22]. Knee joint loading was computed using a validated musculoskeletal model of the rat hindlimb [23] and was used to define the boundary conditions of the FE model. The knee joint functions, as well as forces, stresses, strains, and fluid pressures, were assessed within the femoral and tibial cartilages, and menisci. We suggest that this animal-specific approach could be useful for understanding mechanisms leading to OA progression and may offer valuable insights when evaluating potential treatments in preclinical animal models.

## Materials and Methods

### Magnetic resonance imaging protocol and segmentation

An intact right lower limb of a cadaveric rat without known musculoskeletal disorders (Sprague Dawley, 56-week-old male, body weight = 5.5 N) was immersed in a phosphate buffered saline solution and imaged at room temperature using an 11.74T μMRI scanner in combination with a 10-mm diameter proton RF coil (UltraShield 500 MHz, Bruker BioSpin MRI GmbH, Ettlingen, Germany). MRI was conducted at the facilities of the Kuopio Biomedical Imaging Unit at A.I. Virtanen Institute of Molecular Sciences (University of Eastern Finland, Kuopio, Finland). The MRI data was acquired using ParaVision 6.0.1. software (Bruker) and a 3-D multi-echo gradient echo (MGE) pulse sequence. The imaging parameters were: echo time (TE) = 1.8, 4.9, 8.0, 11.1, 14.2, and 17.3 ms, repetition time (TR) = 100 ms, flip angle (FA) = 20°, field of view (FOV) =14.25 × 9.5 × 9.5 mm^3^, echo spacing (ES) = 3.1 ms, averages = 1, scan time = 1h 49 min, receiver bandwidth = 0.15 MHz and an acquisition matrix of 384 × 256 × 256, yielding an isotropic voxel size of 37μm.

Knee joint geometries that included femoral and tibial cartilages, menisci, collateral, and cruciate ligament insertions were segmented using the open software 3DSlicer (http://www.slicer.org)[24] from the MRI data acquired with the shortest TE. The segmented geometries were imported into Abaqus (v2018; Dassault Systèmes Simulia Corp, Providence, RI) where the FE meshes were constructed using 8-node hexahedral linear poroelastic (C3D8P) elements (Fig 1).

**Fig 1.**
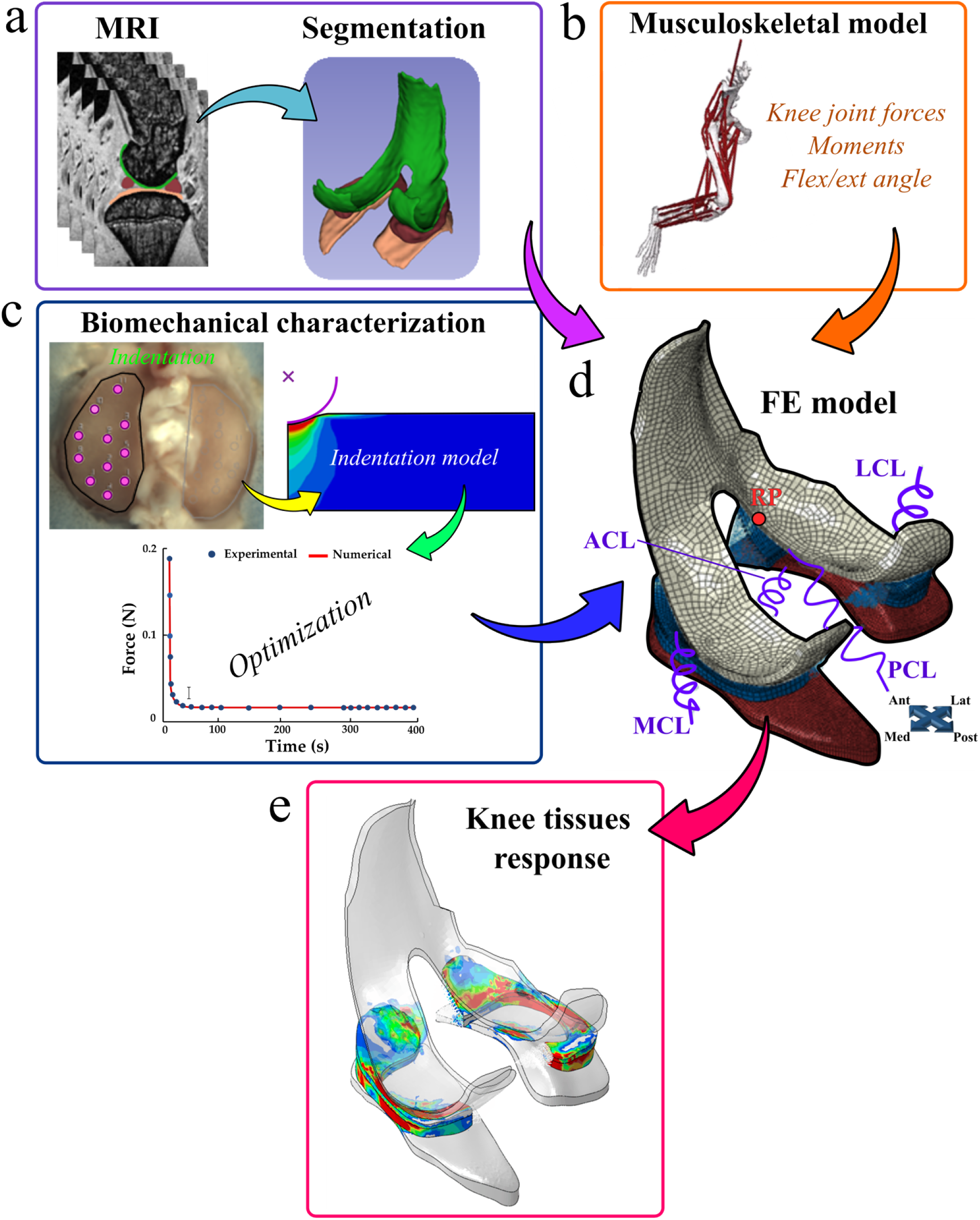
Workflow of the study. (a) Rat knee geometry, (b) motion and loading during gait from a musculoskeletal model, and (c) FRPE material properties from indentation tests were implemented into (d) the FE model. e) Knee tissues’ mechanical responses were evaluated during the stance phase of the gait cycle.

### Musculoskeletal modeling of rat hindlimb

We utilized a previously validated musculoskeletal model of the right hindlimb of Sprague-Dawley rat in OpenSim (SimTK, Stanford, CA) [23,25] (https://simtk.org/projects/rat_hlimb_model). The model was used to determine the knee joint contact forces and lower extremity muscle forces occurring during the gait cycle, which were used as boundary conditions for the FE knee joint model [26]. Briefly, the musculoskeletal model was composed of four body segments, including accurate representations of the bones (spine, femur, tibia, and foot), 14 degrees of freedom, and 39 muscle-tendon actuators that are represented as linear elements in each muscle segment. We prescribed the joint angle profiles during the stance phase of gait by scaling the locomotion and ground reaction force (GRF) data from Charles et al. [27,28] to match the normal (healthy) gait pattern of Sprague-Dawley rats reported in previous experimental studies [29–34]. The scaling of the joint angle-time curves was conducted using a custom MATLAB script (R2019b; The MathWorks, Natick, MA). Scaled joint angles and GRFs were used for estimating the muscle forces using static optimization (e.g. minimizing the cost function associated with muscle activations) and subsequently performed the joint reaction analysis. Finally, the musculoskeletal model outputs: the flexion-extension angle, valgus-varus and internal-external passive moments, and translational knee forces (distal-proximal, medial-lateral, and anterior-posterior) were used to drive the knee joint FE model, by following a similar protocol as previously published [16,35,36] (Fig 2).

**Fig 2.**
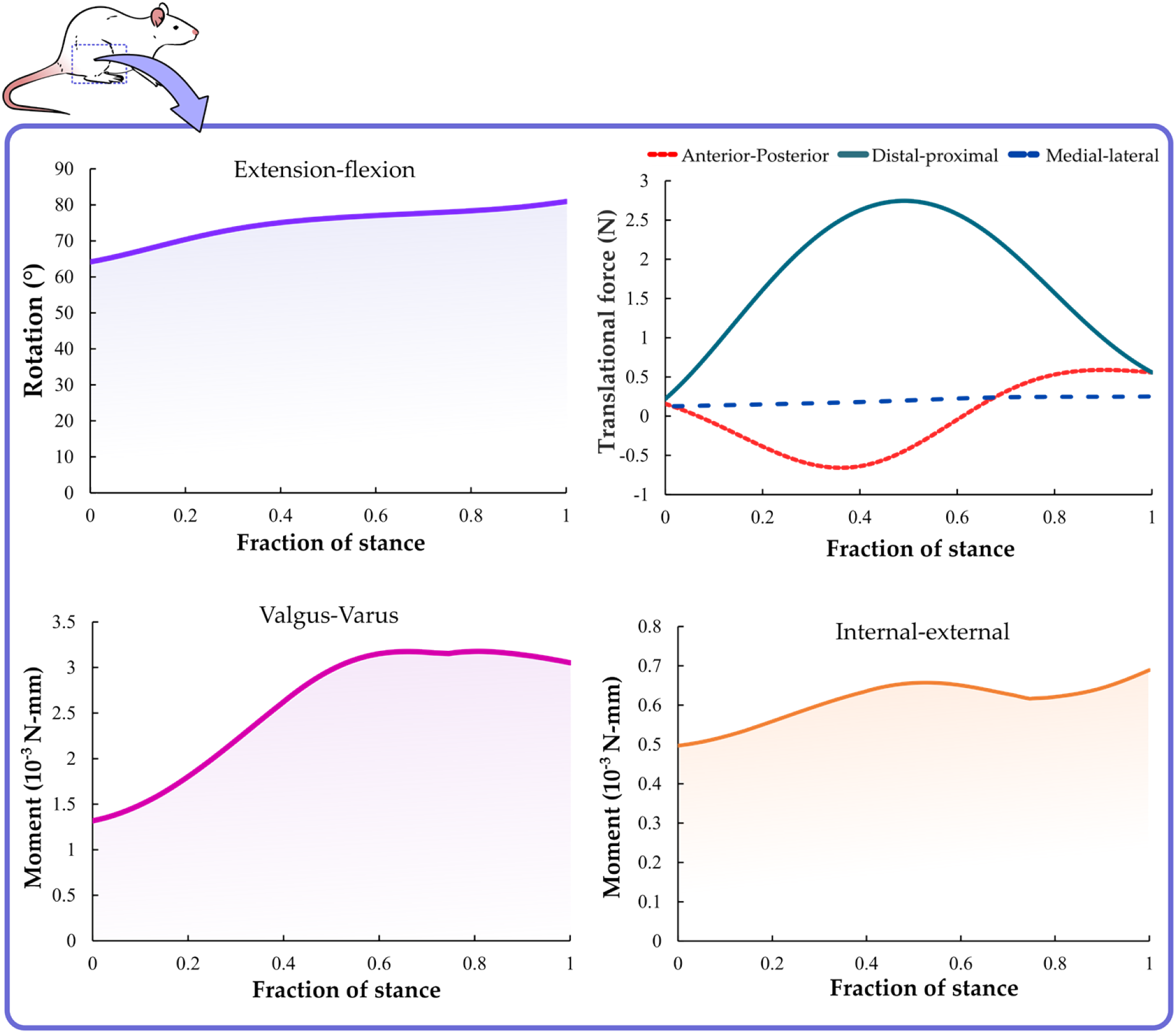
Gait data for the computational model of the rat knee joint. External-internal and valgus-varus moments, and flexion-extension rotation. In addition, anterior-posterior, distal-proximal, and medial-lateral translational forces were implemented in the FE model of the knee joint. The inputs of the FE joint model (joint kinematics and translational forces) were similar to previous experimental studies with Sprague-Dawley rats [29–34].

### Biomechanical articular cartilage characterization

The fibril-reinforced poroelastic (FRPE) properties of healthy Sprague-Dawley rat cartilage were characterized using previously published experimental indentation measurements [22]. The FRPE cartilage parameters were obtained by fitting the stress relaxation curve of the FE model to the mean stress relaxation curve collected from healthy control animals (*n* = 6) of a previous study [22]. Briefly, stress relaxation experiments were performed using a spherical indenter (*r* = 175 ± 2.5 μm, 316 L glass) that was mounted to a multiaxial load cell (force resolution: Fz =3.5 mN and Fx = Fy = 2.5 mN) and a three-axis mechanical tester (Mach-1 v500css, Biomomentum, QC, Canada). For each specimen, the tibial cartilage was fixed in a specimen holder using dental cement and immersed in a phosphate buffered saline solution. To ensure proper sample-indenter contact for consistent and repeatable measurements, an automatic contact criterion of 0.01 N (contact velocity: 0.1 mm/s) was applied to all the samples. Then a single stress-relaxation step (indentation amplitude: 0.04 mm (~30% of uncompressed cartilage thickness), compression velocity: 0.04 mm/s, relaxation time: 400 s) was performed on 11 sites each for the lateral and medial tibial cartilage using the automated indentation mapping system (Fig 1c). After the indentation experiments, the thickness was measured on new 11 sites (located close to those previously identified for the indentation mapping) each for the lateral and medial tibial cartilage using automated thickness mapping with a needle probe.

Subsequently, six axisymmetric FE models of a cylindrical specimen (radius: 1.5 mm) that took into account sample-specific thickness were constructed in Abaqus to simulate the mean of the indentation tests for each sample. The sample height was set to be the mean cartilage thickness measured for each sample (see Table S1 in the **supplementary material**). The geometry was meshed by 825 linear axisymmetric pore pressure continuum elements (element type CAX4P). Mesh convergence was ensured for each model. An FRPE constitutive formulation was implemented for simulating the articular cartilage response [37,38]. Specifically, the material model assumes that cartilage tissue is composed of solid and fluid matrices. The solid matrix is separated into a porous non-fibrillar part, representing the proteoglycan matrix, and an elastic fibrillar network, describing the collagen fibrils. The total stress is given by

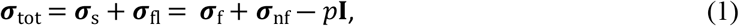

where ***σ***_tot_ is the total stress tensor, ***σ***_s_ and ***σ***_fl_ represent the stress tensors of the solid matrix and interstitial fluid, respectively, *p* is the hydrostatic pressure, **I** is the unit tensor, and ***σ***_f_ and ***σ***_nf_ are the stress tensors of the fibrillar and non-fibrillar matrices, respectively. A neo-Hookean material is utilized to define the non-fibrillar component, in which the stress tensor is given by

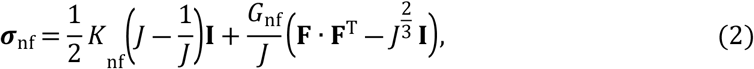

where *K*_nf_ and *G*_nf_ are the bulk and the shear moduli of the non-fibrillar matrix and *J* is the determinant of the deformation gradient tensor **F**. The bulk (*K*_nf_) and shear (*G*_nf_) moduli of the non-fibrillar matrix are established as

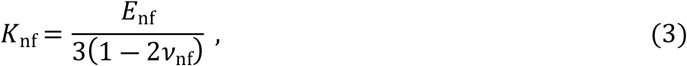

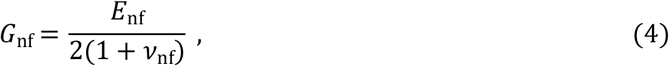

where *E*_nf_ and *ν*_nf_ are the Young’s modulus and the Poisson’s ratio of the non-fibrillar matrix. Then, the stresses in the elastic collagen fibrils are given by

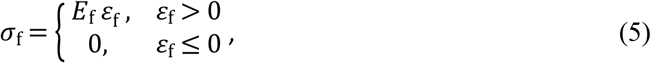

where *σ*_f_ and *ε*_f_ represent the stress and strain of the fibril, and *E*_f_ is the fibril network modulus [38]. Therefore, collagen fibrils support tension only. The fibril network stress emerges from the sum of the primary and secondary collagen fibril stresses, which are computed individually for each integration point in each element [39]. The stresses for these fibrils in tension are defined

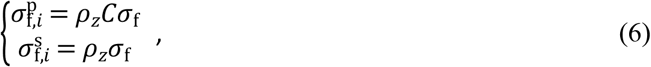

where 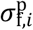 and 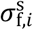 are the stresses for primary and secondary fibrils, respectively, *ρ*_*z*_ is the relative collagen density, and *C* is the density ratio between primary and secondary fibrils. Then, the total stress tensor of the fibrillar network is defined as the sum of the stresses in each fibril (*σ*_f,*i*_):

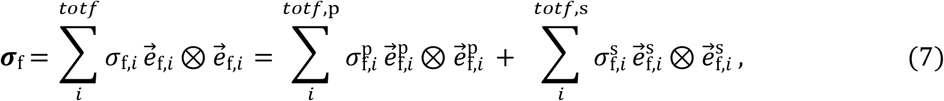

where *totf* is the total number of fibrils, 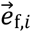 is the fibril orientation vector, *totf*,p and *totf*, s are the total number of primary and secondary fibrils, respectively, and 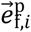 and 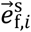 are the primary and secondary fibril orientation unit vectors, and ⊗ represent the outer product. Moreover, the fluid flow in the non-fibrillar matrix is assumed to follow Darcy’s law:

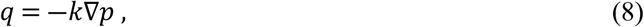

where *q* is the fluid flux in the non-fibrillar matrix, ∇*p* is the hydrostatic pressure gradient vector across the region, and *k* is the hydraulic permeability. The hydraulic permeability is defined to be strain-dependent:

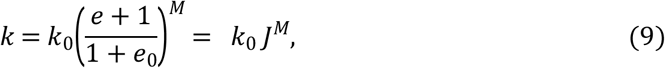

where *k*_0_ is the initial permeability, *M* is a positive constant, and *e* and *e*_0_ are the current and initial void ratios, respectively [39]. The void ratio *e* is expressed by the ratio of the fluid to the solid volumetric fraction:

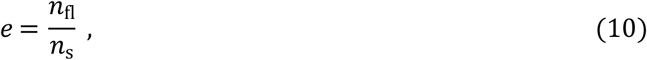

where *n*_s_ is the solid volume fraction and *n*_fl_ is the fluid volume fraction.

The following boundary conditions were used for the axisymmetric FE models, similar to previous reports [37,40]. The bottom of the cartilage sample was fixed in the axial and lateral directions and fluid flow was allowed through the free non-contacting surfaces. However, no fluid flow was allowed to occur at the bottom surface. The contact between the indenter (simulated as a rigid analytical surface) and cartilage surface was assumed impermeable and frictionless. The cartilage sample was subjected to the indentation protocol described earlier in this study. In addition, the FRPE material properties (*E*_f_,*E*_nf_,*k*_0_, and *M*) were obtained by minimizing the normalized mean squared error between the experimentally measured and the FE model-predicted forces using a minimum search algorithm (*fminsearch* function) in combination with Abaqus [37]. Poisson’s ratio of the nonfibrillar matrix was assumed to be 0.42 [40,41], leading to an effective (i.e. apparent) cartilage Poisson’s ratio of ~0.1.

### Finite element model of the rat knee joint

Cartilages and menisci were modeled using the FRPE material. For tibial and femoral cartilages, the fitted FRPE material parameters, depth-dependent collagen fibril architecture, and fluid fraction distribution were implemented [36,42,43]. The tibial cartilage-bone interface was fixed in all directions and bones were assumed rigid. For the menisci, the primary fibrils of the collagen network were oriented circumferentially, and the fluid fraction was assumed to be homogeneously distributed [44–46]. Menisci properties were adopted based on earlier experiments on human meniscus due to a lack of information about rat menisci properties in the literature. In addition, the roots of the menisci were attached to the bone using linear spring elements with a total stiffness of 350 N/mm at each root [47]. A complete list of the material parameters used is given in Table 1.

**Table 1.** Material parameters implemented for cartilage and menisci.

| Material parameter | Cartilage | Menisci |
| --- | --- | --- |
| $E_f$ (MPa) | 3.13<sup>†</sup> | 3.79<sup>□</sup> |
| $E_{nf}$ (MPa) | 0.83<sup>†</sup> | 0.08<sup>*</sup> |
| $k_0$ ($10^{-15} \text{m}^4 \text{N}^{-1} \text{s}^{-1}$) | 3.30<sup>†</sup> | 0.08<sup>*</sup> |
| $\nu$ | 0.42<sup>§</sup> | 0.3<sup>*</sup> |
| $M$ | 1.67<sup>†</sup> | 12.1<sup>*</sup> |
| $C$ | 12.16<sup>**</sup> | 12.16<sup>**</sup> |
| $n_{fl}$ | $0.8 - 0.15h^{**}$ | 0.72<sup>‡</sup> |

*E*_f_ = fibril network modulus, *E*_nf_ = nonfibrillar matrix modulus, *k*_0_= initial permeability, ν = Poisson’s ratio of the nonfibrillar matrix, *M* = exponential term for the strain-dependent permeability, *C* = ratio of primary to secondary collagen fibers, *n*_fl_ = depth-wise fluid fraction distribution, *E*_Ɵ_ = circumferential Young’s modulus of menisci, *n*_f,p_= number of primary fibrils and *h* indicates the normalized distance from the cartilage surface (surface = 0, bottom = 1).

†Obtained from fitting the model to indentation experiments.

^◻^The fibril network modulus of menisci was computed as follows: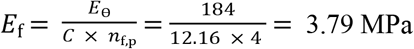

*Danso et al. [44,45]

**Wilson et al. [39]

‡Makris et al. [48]

‡‡Mow and Ratcliffe. [49]

§Korhonen et al., Mäkelä et al. [40,50]

Cruciate ligaments (ACL and PCL) and collateral ligaments (MCL and LCL) were modeled using spring elements with a bilinear behavior. The ligaments were assumed to be pre-elongated (MCL and LCL = 4% [51], ACL and PCL = 5% [36]) of the initial length at the segmented distance by using the bilinear spring selection. The stiffness of the ligaments (MCL 20 N/mm, LCL 20 N/mm, ACL 35 N/mm, and PCL 35 N/mm) were obtained from previous rat ligaments experimental studies [52,53]. The springs were attached to the center of the anatomical attachment sites of each ligament measured from MRI data [36,42]. Ligament anchorage points were fixed at the tibial bone sites during the gait cycle. The anchorage points at the femoral site were coupled to the main reference point (located at the midpoint between the condyles of the femur), allowing them to move along with the rigid bone.

The following boundary conditions were applied to the FE model of the rat knee joint. The stance phase of the rat’s gait obtained from the musculoskeletal model was implemented to drive the FE simulation, similarly to that described in human knee joint studies [36,42]. In detail, after an initial contact step, the flexion-extension angle, and joint moments and translational forces during the stance phase were computed and implemented to the main reference point, located at the mid-point between the lateral and medial epicondyles of the femur (Fig 1). Surface-to-node contacts with frictionless sliding properties were applied between the cartilage-cartilage and cartilage-meniscus contact surfaces. The average and maximum tissue mechanical responses, including maximum principal stress, maximum principal strain, and fluid pressure were analyzed in the knee joint during the entire stance phase of the gait cycle. For evaluating the average tissue responses, average values over the cartilage-cartilage contact area were computed as a function of time.

## Results

### FRPE characterization of articular cartilage

The FRPE material model successfully described the response obtained from the indentation experiments, revealing *R*^2^ = 0.97 ± 0.03 for the coefficient of determination. The optimized FRPE parameters *E*_f_, *E*_nf_, *k*_0_, and *M* (mean ± standard deviation) were 3.13 ± 2.56 MPa, 0.83 ± 0.21 MPa, 3.30 ± 3.00 × 10^−15^ m^4^N^−1^s^−1^, and 1.67 ± 0.62, respectively. Subsequently, the mean value of each optimized cartilage parameter was used for the FE knee joint model (Table 1).

### Finite element model of the rat knee joint

The FE rat knee joint model showed that the maximum principal stress was concentrated on a small area at the beginning of the stance phase (Fig 3). Total tibiofemoral reaction forces obtained in the medial and lateral compartments are presented in Figs. 4a and 4b, respectively. The model calculated the highest tibiofemoral reaction forces (1.16 BW) at ~55% of the stance phase. Furthermore, the secondary knee kinematics displayed an increase in the posterior-anterior and medial-lateral translations at the end of the stance phase (Figs 4c-d). In contrast, the inferior-superior translation decreased with time during stance (Fig 4e). Additionally, the valgus-varus and external-internal rotations increased with stance time (Figs 4f-g).

**Fig 3.**
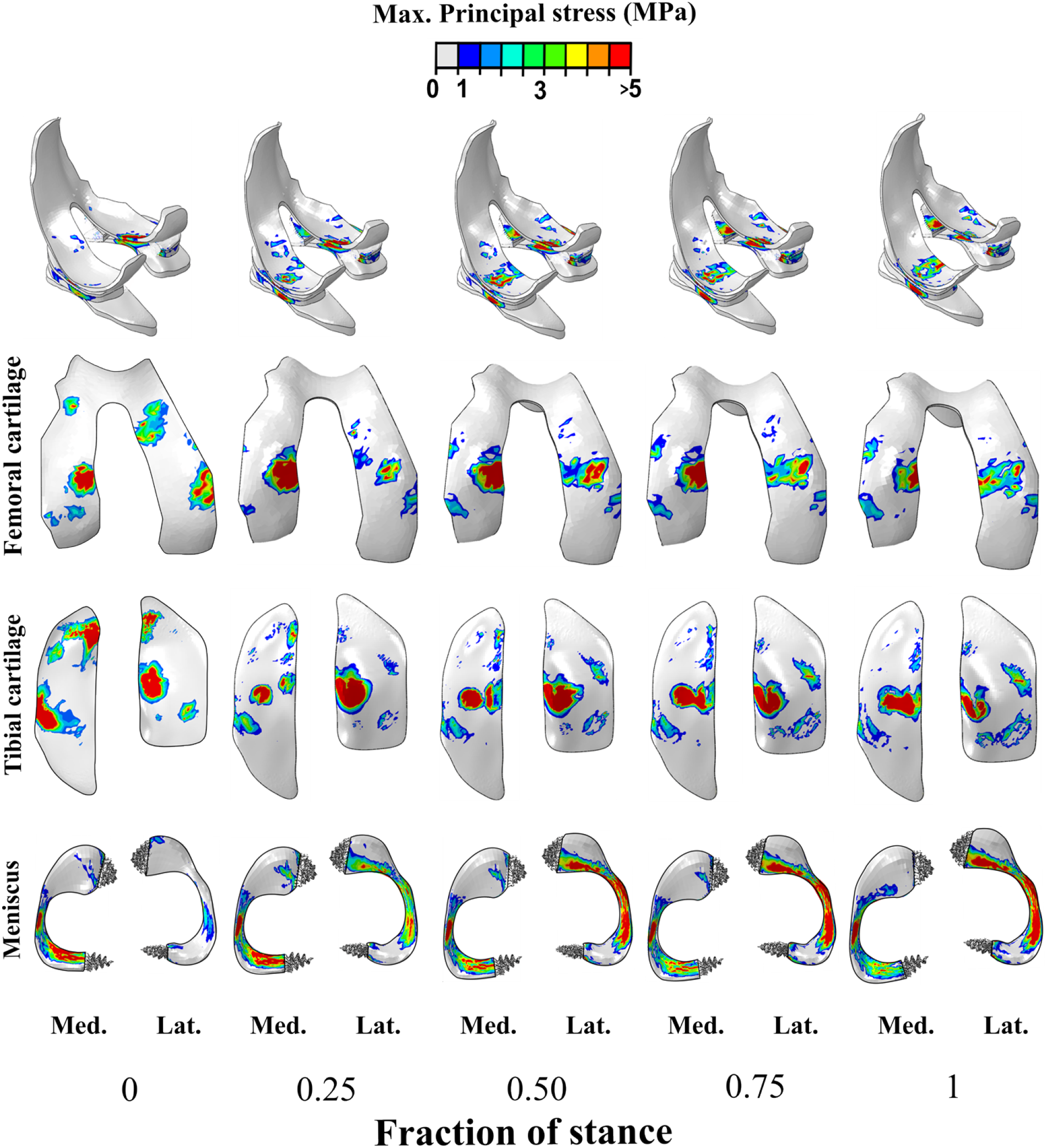
Maximum principal stress distribution in the femoral and tibial cartilages, and menisci calculated from the FE model of the knee joint during the stance phase of the gait cycle (Lat: lateral: Med: medial). The cartilage stresses obtained from the FE model agree with previous numerical studies on mice knee joints under axial compressive forces [18,21].

**Fig 4.**
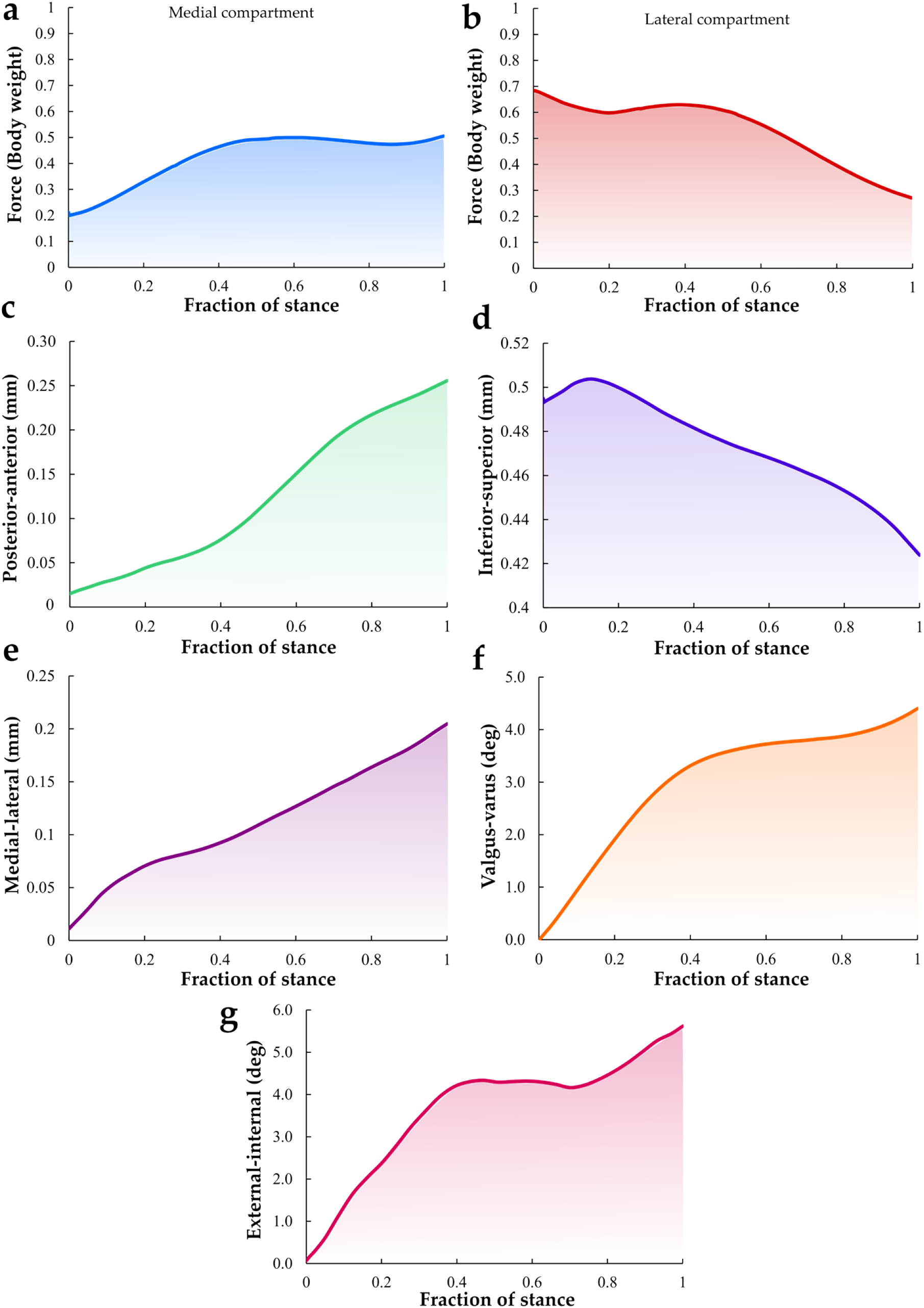
Total tibiofemoral joint reaction force in the (a) medial and (b) lateral compartments, respectively. Translations (c-e) and rotations (f, g) of the tibia with respect to the femur during the stance phase of gait were also presented.

Quantitative analysis of the average tissue mechanical responses over the cartilage-cartilage contact area within the medial and lateral compartments of the tibial cartilage during the stance phase of gait is presented in Fig 5. The average contact pressure and maximum principal strain were virtually constant in the medial compartment (0.02 MPa and 10.0%, respectively) whereas, in the lateral compartment, the average contact pressure and maximum principal strain showed the highest values (0.06 MPa and 30%, respectively) at the start of the stance phase and subsequently contact pressure and principal strain decreased with time (Figs 5a-d). Moreover, the average maximum principal stress and fluid pressure within the medial compartment were highest at midstance (5.6 and 4.8 MPa, respectively). In contrast, the average maximum principal stress and fluid pressure in the lateral compartment decreased with time. Similar to the contact pressure and maximum principal strain response, the highest stress and fluid pressure in the lateral compartment occurred at the beginning of the stance phase. Peak contact pressures, stresses, strains, and fluid pressures within the medial and lateral compartment as a function of stance are presented in Fig S1 (See the **supplementary material**).

**Fig 5.**
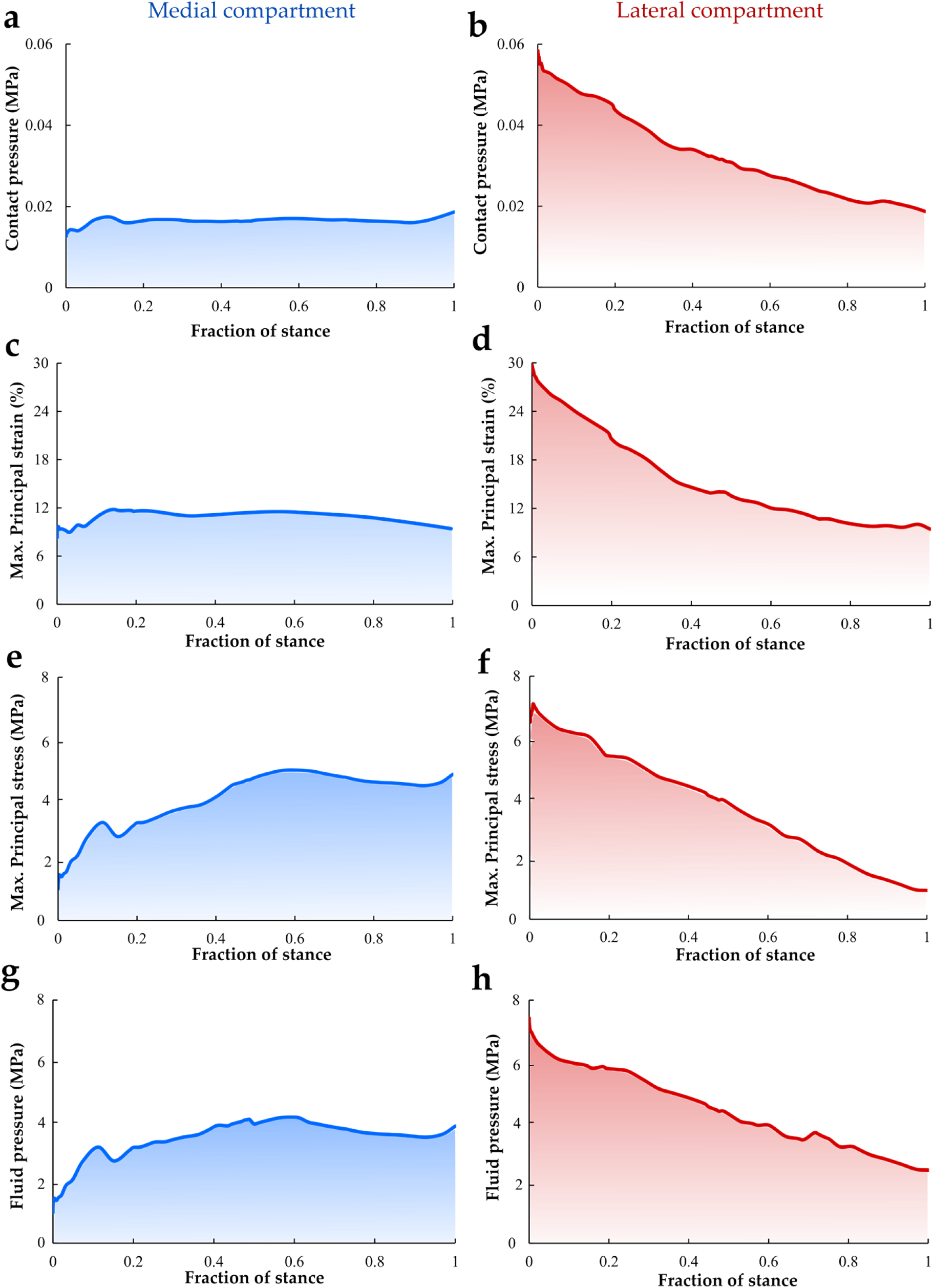
Average contact pressure, maximum principal strain, maximum principal stress, and fluid pressure in the contact area of the medial (a, c, e, and g) and lateral (b, d, f, and h) tibial cartilage surfaces during the stance phase of gait. The contact stresses were similar to a previous study published by Gardner-Morde et al. [20] where average contact stresses in the medial and lateral compartment at the reference loading state were 0.4 and 0.1 MPa, respectively.

The cruciate ligaments (ACL and PCL) in the knee joint experienced higher loads than the collateral ligaments (MCL and LCL) throughout the stance phase (Fig 6). The force in the ACL was higher than that in the PCL. The ACL force decreased from 0 to 30% of the stance phase and then increased until the end of the stance phase (peak ACL load: 3.8 N). In contrast, the PCL force increased from 0 to ~40% of the stance but then decreased until the end of the stance phase (peak PCL load: 2.1 N). Similar to the ACL response, the MCL force (peak MCL load: ~1 N) decreased steadily from the beginning of the stance phase and became virtually unloaded at midstance but increased slightly in the second half of the stance phase. Conversely, the LCL force decreased at the start of the stance phase but revealed a minor increase from ~20% of stance until the end of the gait cycle (peak LCL load: ~0.6 N).

**Fig 6.**
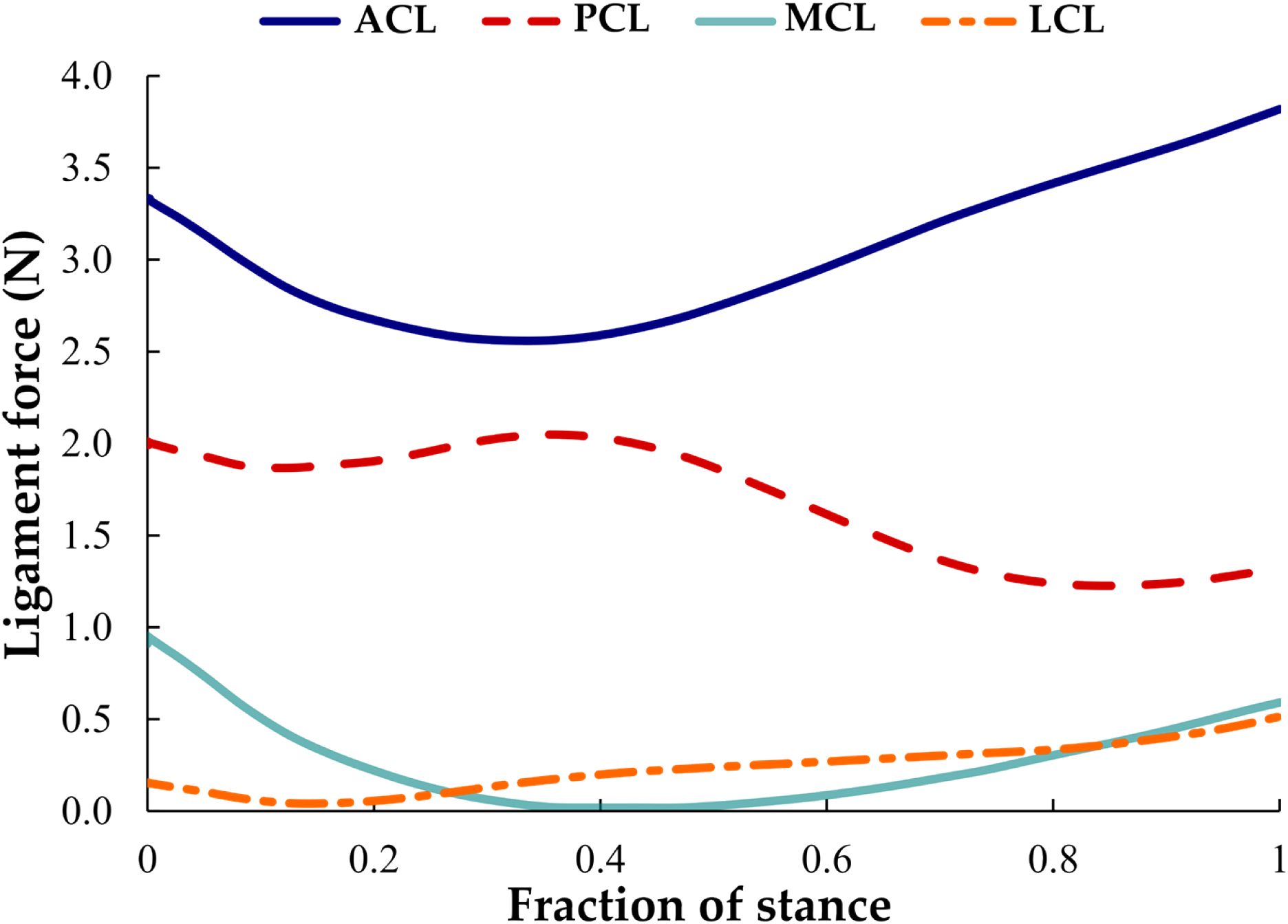
Forces transmitted through the cruciate ligaments (ACL and PCL) and collateral ligaments (MCL and LCL) of the knee joint during the stance phase of gait.

## Discussion

### Summary

In the present study, we described a workflow for the generation and simulation of a finite element model of a rat knee joint to estimate the biomechanical responses of articular cartilage and other knee joint tissues during the stance phase of walking. To the best of our knowledge, this approach represents the first 3-D rat knee model that can be used to investigate the cartilage and meniscus stresses and strains from dynamic joint loading during gait. The rat knee joint geometries were extracted from a 3-D MRI dataset and the boundary conditions regarding loading of the joint were extracted from a musculoskeletal model of the rat hindlimb. In addition, the FRPE properties of the rat cartilage were derived from data of mechanical indentation testing across the articular surfaces of the rat knee. Our numerical results showed the effect of simulating anatomical and locomotion characteristics on the rat knee joint for estimating tissue responses, such as contact pressures, stresses, strains, and fluid pressures, which can be used to estimate mechanobiological changes of tissues during OA as well as to evaluate the effect of knee joint disorders and gait impairments on articular cartilage in preclinical models of joint injury and disease [54].

### Biomechanical evaluation of articular cartilage

Fibril-reinforced poroelastic (FRPE) properties of cartilage in the rat knee joint were obtained using indentation experiments in the tibial plateau, combined with FE models and an optimization algorithm. Although previous studies have measured cartilage poroelastic properties during creep experiments in rat tibial cartilage [55], femoral cartilage [56,57], and mouse tibial plateau [58], our study constitutes the first investigation to describe effectively the mechanical behavior of cartilage of rat knee by using the mechanical moduli of the collagen fibril network and the non-fibrillar solid matrix. In a previous study, Athanasiou et al. [56] performed indentation experiments on rat articular cartilage. The aggregate compressive modulus (comparable to the nonfibrillar matrix modulus *E*_nf_) and permeability of normal/healthy cartilage were 0.75 ± 0.16 MPa and 3.13 ± 2.59 × 10^−15^ m^4^N^−1^s^−1^, respectively. These findings are in agreement with the results of the present study, in which only healthy rat cartilage tissue was used.

Our FE rat knee joint model was able to describe the stress, strain, and contact pressure distributions on cartilage and menisci during the stance phase of gait. Numerical results revealed that the average cartilage contact pressure remained almost constant in the medial compartment, but in the lateral compartment the average contact pressure varied during the gait cycle. In a previous study, Gardner-Morde et al. [20] estimated compressive contact stresses using discrete element analysis of the rat tibiofemoral joint during standing with different applied varus loads without menisci. Average contact stresses in the medial and lateral compartment at the reference loading state were 0.4 and 0.1 MPa, respectively. These contact stresses were of similar magnitude to those reported in this study, in which the meniscus was included in the simulation (0.1 MPa peak contact pressure).

On the other hand, the stress distributions indicated that the medial compartment experienced an increase in the maximum principal stress during gait, while the lateral tibial compartment revealed decreasing values during the second half of the stance phase. The stress distributions and forces indicated that the meniscus provides substantial mechanical support during dynamic gait loading. The magnitude of the cartilage stresses obtained from the FE model agrees with computational studies on mice knee joints under axial compressive forces [18,21].

Regarding the notable mechanical support provided by the meniscus during gait loading, cartilage-meniscus force represented 36% and 42% of the total reaction force within the medial and lateral compartments in the midstance phase of walking (See Fig S2 in the **supplementary material**). This finding is in good agreement with previous observations in mouse FE knee models under weight-bearing conditions, where the cartilage-cartilage contact was reduced by 34% in the presence of the meniscus on the lateral condyle [19,21]. Potentially, our numerical model could elucidate the mechanisms behind the progressive structural changes observed in DMM surgical instability pre-clinical models of OA [59]. Also, for investigating the effect of refining surgical small rodent models of OA on both joint pathology and pain response [60].

ACL and PCL forces were the largest knee joint ligament forces throughout the entire stance phase of gait. This finding supports previous observations that these ligaments are the main joint stabilizers, controlling the anterior-posterior translation of the tibia [61,62]. It is known that ACL and PCL deficiency has an influence on knee joint kinematics and kinetics, increasing the stress concentration in certain areas of the articular cartilage and leading to cartilage degeneration [63–65]. In fact, preclinical posttraumatic OA animal models following ACL rupture have been widely developed [66–68]. Potentially, our current numerical approach can be used to investigate the progression of OA following ACL transection by considering the effect of gait impairments and weight-bearing alterations on the function of the rat knee joint and subsequent changes in the cartilage tissue [5,69,70].

### Limitations

Our study contains limitations regarding the FE model development, animal gait motion, tissue mechanical characterization, and specific assumptions. First, knee tissue geometries were based on a single male Sprague Dawley rat. This single joint might not represent all anatomical details of the rat knee across animals and rat strains, but it is practical for this proof-of-concept study. In the future, a large number of animals should be studied to consider different anatomical characteristics of articular cartilage as a function of age, sex, diet, etc. Second, the gait motion used to drive the FE knee model was extracted from a previous validated musculoskeletal model in combination with a generic locomotion pattern of Sprague-Dawley rats reported in the literature [29–34]. This approach might not completely represent all the hindlimb motion details of an animal during a full gait cycle. However, using generic locomotion data from the literature was sufficient for the methodological development required in this study. In the future, we plan to obtain animal-specific motion using 3-D X-Ray Reconstruction of Moving Morphology (XROMM) [71] in combination with musculoskeletal modeling to acquire the ground reaction forces, moments, and accurate and subject-specific hindlimb kinematics of rats. Third, we did not consider the patella and tendons in the FE model. This might represent differences in the rotations and joint reaction force, but we would not expect greater variations of cartilage stresses and strains than observed in this simpler model. Our workflow could be applied to generate complex models with additional anatomical features, such as the kneecap, tendons, and muscles. We acknowledge that these additional aspects could be included but the animal-specific motion and a more sophisticated musculoskeletal model are necessary to validate the above-mentioned details. Fourth, the characterization of the biomechanical properties of rat cartilage considered only a single stress-relaxation step during a single indentation experiment in tibial cartilage. It was assumed that the femoral cartilage had the same properties as the tibial cartilage. More stress-relaxation steps should be performed to characterize the intrinsic nonlinearities of cartilage tissue across all joint surfaces, and additional mechanical testing (e.g. shear, tension, unconfined compression, and confined compression) should be done to complement the currently available cartilage responses. Fifth, in order to overcome the lack of information on the material properties of the menisci in the rat knee joint, we used values from previous experimental studies [44,72]. Characterization of rat meniscus material properties and implementation of these properties into FE models are part of our upcoming plan. Although experimental tests on rat ligaments and tendons are challenging to conduct due to the small size of the samples, it is worth characterizing both the nonlinear toe and linear regions of the ligaments for a better understanding of the function of ligaments and tendons in healthy, injured, and diseased knees [73]. Sixth, volumetric information of the healthy tibiofemoral joint of the rat was obtained using the presented MRI acquisition scheme with a high isotropic resolution of 37 μm (using an 11.74T μMRI scanner). As the femoral and tibial cartilage thickness is approximately 180 μm, the resolution allowed for four to five pixels across the cartilage thickness, which may affect the accuracy of tissue detection. Partial volume artifacts could further affect the segmentation, but the high resolution utilized helps mitigate this effect. Previous research [74,75] was performed using anisotropic pixels with pixel sizes greater (59 × 117 × 234 μm^3^ and 51 × 51 × 94 μm^3^, respectively) than those used in our work. In addition, chemical shift of the fat, emphasized by the ultra-high field of the magnet (11.74T), may cause errors in the estimation of tissue volumes. The studies mentioned above utilized fat suppression methods in their gradient echo acquisition schemes [74,75]. Here, fat suppression was not used in the main acquisition to preserve as much signal as possible, as preliminary images with fat suppression suggested minimal effect.

#### Future developments

Our model of the rat knee provides a potential numerical tool to estimate the loading and changes in articular cartilage and other tissues of the rat knee during the stance phase of gait after pre-clinical OA interventions in rodents. Cartilage tissue mechanical responses, such as stress, strain, and fluid velocity have been reported as indicators of tissue adaptation and degradation after joint injury and/or disease [16,76,77]. For example, our knee model allows for simulating the effects of ACL transection and partial/total meniscectomy on the compositional and structural changes in cartilage based on mechanobiological response. In this context, the FE models can be used to investigate the effects of interventions in animal models and to estimate adaptations in mechanical properties of knee joint tissues. Furthermore, our numerical model could be used to study the effects of exercise and prebiotic supplementation, described in OA animal models of diet-induced obesity [11,12,78]. For instance, we could combine the body weight, locomotion, and structural properties with cartilage degenerative algorithms in our FE model for predicting OA progression [76,79].

Longitudinal observations of OA progression have been conducted using quantitative μMRI in the knee joints of rats subjected to different interventions [80,81]. These cartilage properties obtained from MRI could be included in the FE model for evaluating the structural progression of OA as well as for validating the numerical predictions driven by different degenerative mechanisms.

## Conclusions

We present a workflow for the generation and simulation of FE models of the rat knee joint. Our model considers both the anatomical and locomotion characteristics of the rat knee joint for estimating tissue mechanical responses in the articular cartilage. In the future, we will expand this approach to investigate tissue adaptations based on the mechanobiological response of the cartilage tissue to controlled interventions. Thus, our numerical FE model employing FRPE material properties may allow for studying the mechanisms leading to changes in composition and structure in cartilage after a traumatic injury or specific pre-clinical interventions. After these evaluations and further validation of the numerical predictions, this model could be applied in the planning of joint loading to prevent the progression of knee joint OA.

## Acknowledgments

The authors appreciate the support of the University of Eastern Finland, Lund University, and the University of Calgary to undertake this study.

## Author contributions

**Conceptualization:** GAO, KK, EKM, LS, PT, JLR, TVT, MJN, HI, WH, RKK

**Data curation:** GAO, KK, LS, PT, TVT, MJN

**Formal analysis:** GAO, KK, LS, TVT, MJN

**Funding acquisition:** PT, MJN, HI, WH, RKK

**Investigation:** GAO, KK, EKM, LS, PT, JLR, TVT, MJN, HI, WH, RKK

**Methodology:** GAO, KK, EKM, LS, PT, JLR, TVT, MJN, HI, WH, RKK

**Project administration:** HI, WH, RKK

**Resources:** HI, WH, RKK

**Software:** GAO, KK

**Supervision:** EKM, LS, PT, MJN, HI, WH, RKK

**Validation:** GAO

**Visualization:** GAO

**Writing– original draft:** GAO

**Writing–review & editing:** GAO, KK, EKM, LS, PT, JLR, TVT, MJN, HI, WH, RKK

## Data availability

The datasets generated and analyzed during this study are available from the corresponding author on a reasonable request.

## Compliance with Ethical Standards

### Funding

We acknowledge funding support from the European Union’s Horizon 2020 research and innovation programme under the Marie Skłodowska-Curie grant agreement No 713645, Strategic Funding of the University of Eastern Finland, Academy of Finland (grant nos. 285909, 324529, 324994, 328920, 332915, 334773 - under the frame of ERA PerMed), Swedish Research Council (2019-00953 - under the frame of ERA PerMed), European Regional Development Fund, Innovation Fund Denmark (9088-00006B - under the frame of ERA PerMed), Finnish Cultural Foundation (grant #191044 and #200059), Maire Lisko Foundation, Sigrid Juselius Foundation, Päivikki ja Sakari Sohlberg Foundation, Maud Kuistila Memorial Foundation, Saastamoinen Foundation, The Killam Foundation, The Canada Research Chair Programme and The Canadian Institutes for Health Research.

